# Selective accumulation of matrix proteins inside of peroxisomal subdomains

**DOI:** 10.1101/2022.10.02.510529

**Authors:** Julia Ast, Nils Bäcker, Domenica Martorana, Christian Renicke, Thorsten Stehlik, Thomas Heimerl, Christof Taxis, Gert Bange, Michael Bölker, Björn Sandrock, Kay Oliver Schink, Johannes Freitag

## Abstract

Formation of specialized reaction spaces prevents interference between distinct cellular pathways. Peroxisomes are cellular compartments involved in a large diversity of metabolic processes. How peroxisomes differentiate into subpopulations and by which mechanism intraorganellar domains are formed remains largely elusive. Here, we report on enzymes from the fungus *Ustilago maydis*, which accumulate inside of peroxisomal subdomains. We describe a short peptide motif (Thr-Ile-Ile-Val) sufficient to trigger focal localization. Mining for proteins with similar motifs uncovered several peroxisomal matrix proteins that accumulate in intraorganellar foci. These foci are enriched in the enzyme urate oxidase – a typical constituent of the paracrystalline core of peroxisomes. Upon peroxisome proliferation uneven distribution of focal structures results in the formation of peroxisome subpopulations with different protein content. The underlying principle of subdomain formation is evolutionary conserved in mammalian peroxisomes and formation of similar foci was also observed inside of mitochondria. We propose that peroxisomal proteins show an individual propensity to self-assemble. This formation of protein aggregates appears to be a ubiquitous driving force to spatially organize the peroxisomal proteome.

## Introduction

Peroxisomes are cellular compartments surrounded by a single membrane. They contain many metabolic enzymes, in particular those required for degradation of fatty acids and detoxification of hydrogen peroxide (Poirier et al., 2006; Smith and Aitchison, 2013). In humans, peroxisomes are essential and mutation of peroxisomal biogenesis proteins (Peroxins or Pex proteins) leads to severe disorders including the Zellweger syndrome (Wanders, 2014). Import of proteins into the matrix of peroxisomes requires either of two cytosolic targeting factors, Pex5 and Pex7 (Francisco et al., 2017; Kim and Hettema, 2015). They recognize specific sequence motifs located either at the Carboxy (*C*)-terminal end of the cargo protein (peroxisomal targeting signal type 1; PTS1) or within the Amino (*N*)-terminal part (peroxisomal targeting signal type 2; PTS2) (Lazarow, 2006; Walter and Erdmann, 2019). The prototype PTS1 bound by Pex5 is a tripeptide with the amino acid sequence Ser-Lys-Leu (Gould et al., 1989).

Fungal peroxisomes are diverse and contain specialized metabolic pathways e.g. for secondary metabolite and siderophore biosynthesis, methanol degradation and carbohydrate metabolism (Bartoszewska et al., 2011; Freitag et al., 2012; Gründlinger et al., 2013; Stehlik et al., 2014; van der Klei and Veenhuis, 2006). A peculiar variant of peroxisomes identified in *Neurospora crassa* are Woronin bodies, which seal septal pores in wounded hypha (Jedd and Chua, 2000; Nguyen et al., 2021). Differentiation of peroxisomes into Woronin bodies is achieved through expression of the Hex1 protein, which forms large crystals in Woronin bodies and is their major constituent (Pieuchot and Jedd, 2012; Tey et al., 2005).

The mechanisms by which peroxisomes differentiate into subpopulations in other biological systems or how they form intraorganellar domains are largely uncharacterized. Isolation and biochemical characterization of rat liver peroxisomes revealed peroxisomal populations with different protein content (Völkl et al., 1999). In yeast, distinct peroxisome populations can be distinguished by means of their age (Kumar et al., 2018). In plants, different types of peroxisomes emerge during development (Hu et al., 2012; Lingard et al., 2009). High-resolution imaging studies uncovered the presence of subdomains inside of the peroxisomal membrane of the yeast *Saccharomyces cerevisiae* (Galiani et al., 2016). Early electron microscopy and biochemical studies demonstrated that the enzyme urate oxidase is a major constituent of an electron-dense paracrystalline core structure (Rouiller & Bernard, 1956; De Duve, 1960; Baudhuin et al, 1965; De Duve & Baudhuin, 1966; Völkl et al, 1988).

Here we show, that a short peptide motif drives the accumulation of peroxisomal matrix proteins into subdomains, which we propose to be related to the electron-dense paracrystalline core. Focal accumulation determines both the protein distribution within single peroxisomes and the formation of peroxisomes with different protein content upon organelle proliferation.

## Results

### Identification of peroxisome subpopulations in *U. maydis*

We have previously shown that in *U. maydis* the acyltransferases, Mac1 and Mac3, involved in biosynthesis of surface-active mannosylerythritol lipids (MELs), are localized to peroxisomes (Becker et al., 2021; Bölker et al., 2008; Freitag et al., 2014; Kämper et al., 2006). Upon a closer inspection of fluorescence microscopic images we noticed that both enzymes are not homogenously distributed among the entire population of peroxisomes (Fig. 1A).

**Figure 1.**
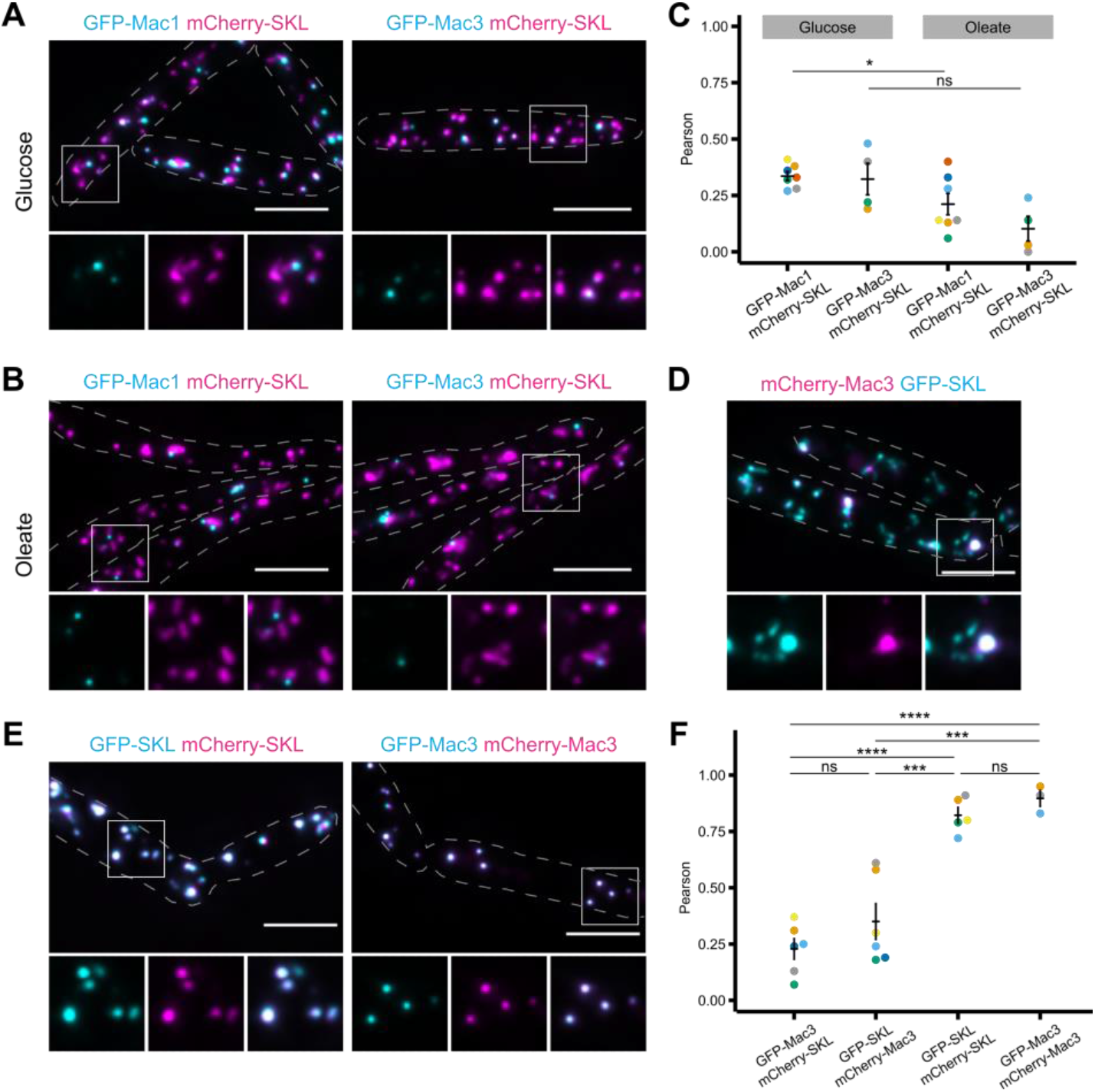
Acyltransferases Mac1 and Mac3 accumulate in peroxisome subpopulations. *U. maydis* strains expressing N-terminally GFP-tagged versions of Mac1 and Mac3 (cyan) and the peroxisomal marker protein mCherry-SKL (magenta) were inspected by epifluorescence microscopy. Representative images of glucose grown cells (A) or oleate grown cells (B). Full images are shown as overlays of the green and red channel. For insets single channels and merged channels are depicted. (C) Quantifications show Pearson’s correlation coefficients of GFP and mCherry signals for indicated strains. (D) Representative epifluorescence image of cells expressing mCherry-Mac3 together with GFP-SKL organized as described above. (E) GFP-SKL and mCherry-SKL or GFP-Mac3 and mCherry-Mac3 were co-expressed and representative images are shown organized as described above. (F) Quantification show Pearson’s correlation coefficients of GFP and mCherry signals for indicated strains. (A); (B); (D) and (E) Scale bars: 5 μm.

Green fluorescent protein (GFP) tagged Mac1 or Mac3 showed only partial colocalization with the red fluorescent peroxisomal reporter mCherry-SKL (Fig. 1A). Oleic acid is known to induce peroxisome proliferation in various species including *U. maydis* (Camões et al., 2015; Kohlwein et al., 2013). Four hours of incubation in oleic acid-containing medium substantially enhanced the observed subpopulation phenotype (Fig. 1B). Quantifications revealed low correlation coefficients for the red and the green signal (Figs. 1C, S1A and S1B). Reciprocal exchange of fluorescent tags did not alter our results indicating that it was not the specific fluorescent protein, which caused the remarkable enrichment in peroxisome subpopulations (Fig. 1D). As a control, we expressed either the combination of GFP-Mac3 and mCherry-Mac3 or the combination of GFP-SKL and mCherry-SKL and observed a high degree of co-localization (Fig. 1E). In aggregate these data show that both acyltransferases are localized only in a fraction of peroxisomes suggesting that in *U. maydis* specific populations of peroxisomes with different protein content exist.

### Accumulation of Mac1 and Mac3 inside of a subdomain of peroxisomes

GFP-Mac1 or GFP-Mac3 fluorescence seemed not to fill the entire peroxisome but was enriched in distinct foci that merged with only a section of the organelle stained by mCherry-SKL (e.g. Fig. 1B). Therefore, we analyzed suborganellar localization of GFP-Mac1 and GFP-Mac3 by super-resolution imaging using structured illumination microscopy (SIM) (Gustafsson, 2000). Analysis of SIM images revealed that both fusion proteins indeed concentrate in subdomains of peroxisomes, while mCherry-SKL and GFP-SKL were distributed among the whole organelle (Fig. 2A to 2C). This was as well visualized by 3D-reconstructions (Fig. 2D). Time-lapse imaging showed that foci containing acyltransferases are stable over time (Fig. S2A and Movie 1).

**Figure 2.**
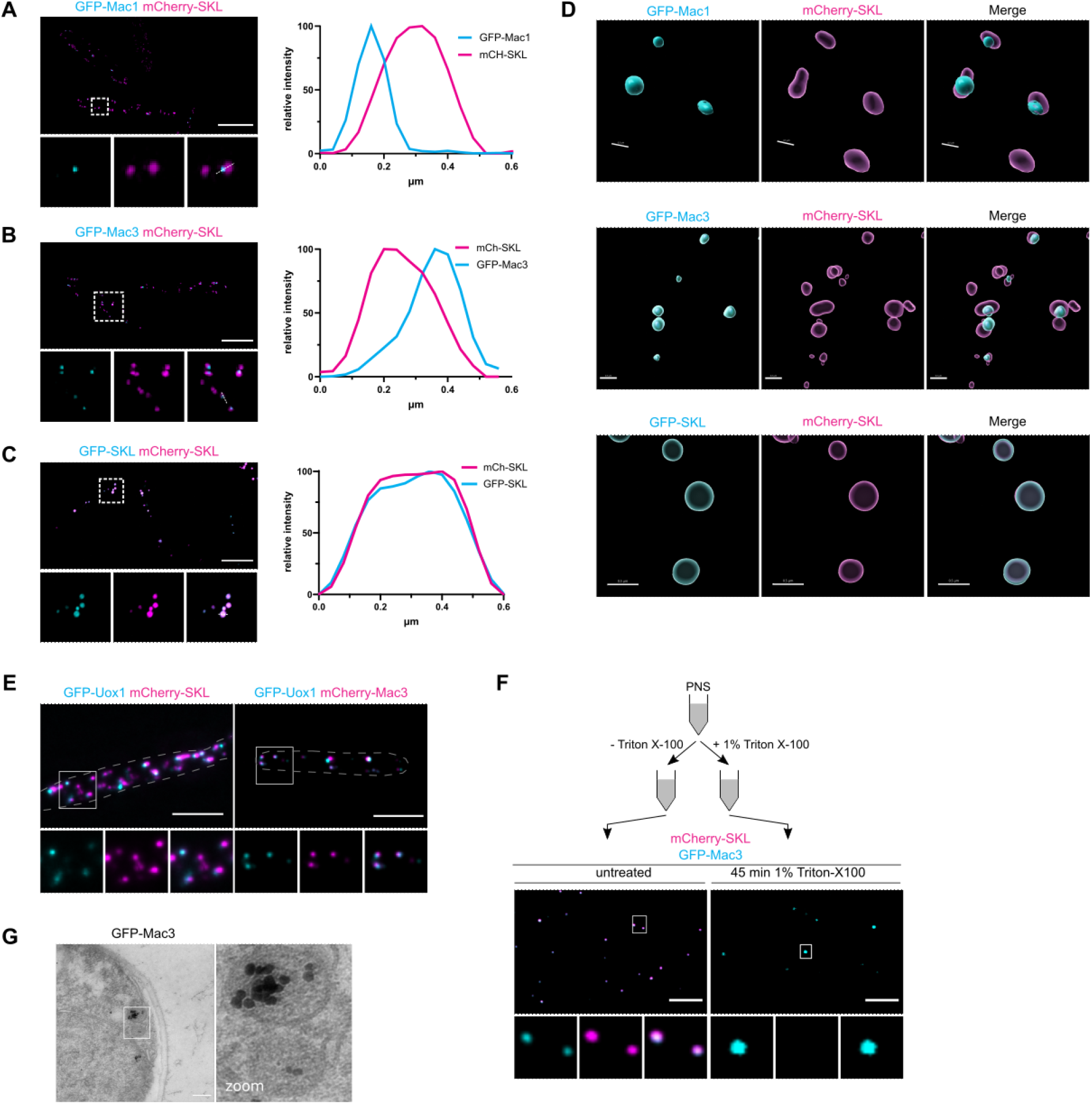
Acyltransferases Mac1 and Mac3 accumulate in peroxisome subdomains also enriched for urate oxidase Uox1. Cells were analyzed by super-resolution microscopy using SIM. Representative images of cells expressing GFP-Mac1 and mCherry-SKL (A), GFP-Mac3 and mCherry-SKL (B) and GFP-SKL and mCherry-SKL (C). Full images are shown as overlays of the green and red channel. For insets single channels and merged channels are depicted. Associated graphs show the normalized fluorescence intensity of GFP and mCherry along the indicated lines. (D) 3D-reconstruction of GFP-Mac3 containing peroxisomal subdomains (cyan). Scale bar: 0.5 μm (E) Representative epifluorescence images of cells expressing GFP-Uox1 with mCherry-SKL (left) or mCherry-Mac3 (right). Organization of pictures as described above. Scale bars: 5 μm. (F) Crude organelle preparations of indicated strains were imaged by epifluorescence microscopy after incubation in lysis buffer supplemented with Triton X-100 (right). Preparations incubated in lysis buffer without Triton X-100 served as control (left). Organization of pictures as described above. Scale bars: 5 μm. (A) – (F) Signals of GFP-tagged proteins are shown in cyan and of mCherry-tagged proteins in magenta. (G) Transmission electron micrograph of cells expressing GFP-Mac3. Staining was achieved through labeling of GFP-Mac3 with anti-GFP and subsequent immunogold labeling followed by silver enhancement. Scale bar: 0.2 μm.

Previous studies of mammalian peroxisomes uncovered a paracrystalline core region, which typically contains the enzyme urate oxidase as a major constituent but also harbors additional proteins (De Duve & Baudhuin, 1966; Völkl *et al*, 1988). We determined the localization of the putative *U. maydis* urate oxidase ortholog (UMAG_00672) fused to GFP (GFP-Uox1). GFP-Uox1 accumulated in foci similar to GFP-Mac1 and GFP-Mac3 and also occurred only in a subpopulation of peroxisomes (Fig. 2E) suggesting that all three proteins are enriched in structures probably representing the paracrystalline core of peroxisomes. Indeed, co-expression of GFP-Uox1 and mCherry-Mac1 or GFP-Uox1 and mCherry-Mac3 resulted in a high degree of colocalization (Fig. 2E and Fig. S2B).

It was reported previously that the paracrystalline core of peroxisomes can sustain treatment with nonionic detergent (Tsukada et al., 1966). We prepared crude organelle extracts from strains expressing mCherry-SKL together with GFP-Mac3, mCherry-Mac3 together with GFP-Uox1 or mCherry-Mac1 together with GFP-Mac3. Triton X-100-treated or non-treated samples were inspected by epifluorescence microscopy. While detection of mCherry-SKL was highly sensitive to treatment with detergent, fluorescently labelled Mac1, Mac3 and Uox1 remained visible inside of detergent resistant punctate structures (Fig. 2F and Fig. S2C). Furthermore, transmission electron microscopy in combination with immunogold labeling showed that GFP-Mac3 is concentrated within a subdomain of peroxisomes (Fig. 2G). Hence, foci enriched for all three enzymes are structures that likely represent the paracrystalline core of *U. maydis* peroxisomes.

### A short peptide motif sufficient for enrichment of proteins inside of the core region

The C-terminal 12 amino acids contribute to targeting efficiency of PTS1 proteins (Brocard and Hartig, 2006). To examine whether information within the peroxisomal targeting signal is involved in formation of subdomains and subpopulations, we analyzed the localization of GFP fused to the C-terminal PTS1 containing dodecamers of Mac1 and Mac3 (GFP-PTS1_Mac1_ and GFP-PTS1_Mac3_) (Fig. 3A). Interestingly, only GFP-PTS1_Mac3_ accumulated in subpopulations and was enriched inside of peroxisomal foci, while GFP-PTS1_Mac1_ colocalized with mCherry-SKL positive peroxisomes and was homogenously distributed within the organelle (Fig. 3B and 3C). By truncation analysis we identified a short peptide motif inside of the 12mer derived from Mac3 consisting of the amino acid stretch Thr-Ile-Ile-Val (TIIV) as required for efficient focal accumulation of GFP (Fig. S3A and S3B). Insertion of this sequence motif between GFP and a canonical SKL containing PTS1 motif induced enrichment of GFP inside of subpopulations of peroxisomes (Fig. 3D). Fusion of the TIIV-tetrapeptide to the C-terminus of a PTS2-GFP reporter protein (Ast et al., 2022) resulted in a similar pattern of localization (Fig. 3E and 3F). This suggests, that focal accumulation does not involve a specific pathway for protein import but probably requires events that occur within the organelle.

**Figure 3.**
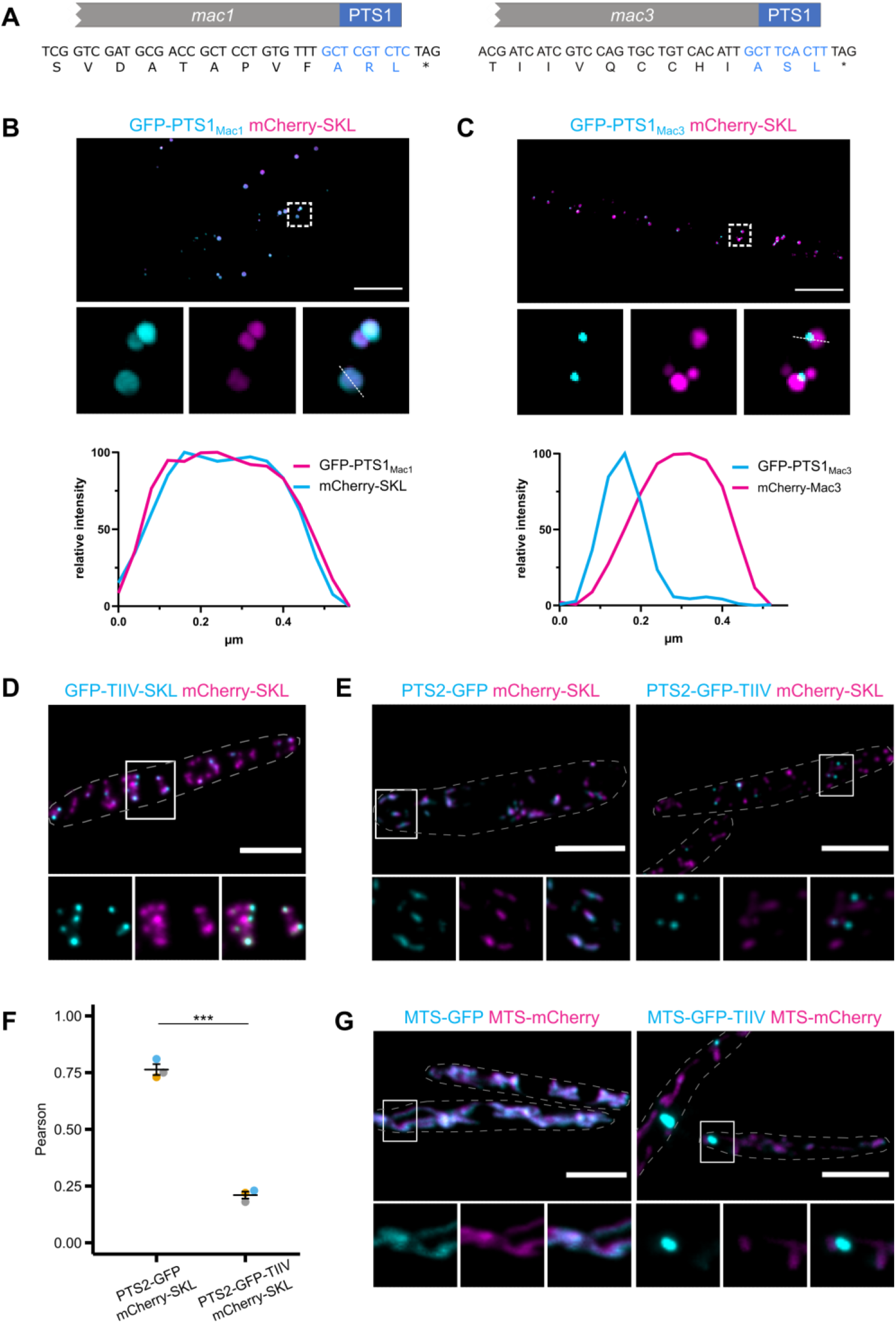
A short peptide motif sufficient for accumulation of proteins inside the core structure. (A) C-terminal dodecamers of Mac1 and Mac3 containing the characteristic PTS1 motifs. Cells were analyzed by super-resolution microscopy using SIM. Representative images of cells expressing GFP-PTS1_Mac1_ and mCherry-SKL (B) or GFP-PTS1_Mac3_ and mCherry-SKL. (C) Full images are shown as overlays of the green and red channel. For insets single channels and merged channels are depicted. Associated graphs show the normalized fluorescence intensity of GFP and mCherry along the indicated lines. (D) GFP fused to a peptide consisting of the small motif Thr-Ile-Ile-Val (TIIV) and the prototype PTS1 Ser-Lys-Leu (SKL) was co-expressed with mCherry-SKL. A representative epifluorescence image is shown. Organization of the picture as above. (E) Cells expressing a PTS2-GFP reporter proteins either with or without a C-terminal TIIV motif were co-expressed with mCherry-SKL. Representative epifluorescence images are shown. Organization of the pictures as above. (F) Quantification show Pearson’s correlation coefficients of GFP and mCherry signals for indicated strains. (G) GFP containing a mitochondrial targeting signal (MTS) either with or without an additional C-terminal TIIV motif was co-expressed with a MTS-mCherry. Representative epifluorescence images are shown. Organization of the pictures as above. ((A) – (E) and (G) Signals of GFP-tagged proteins are shown in cyan and of mCherry-tagged proteins in magenta. Scale bars: 5 μm.

To address if a TIIV motif does provoke subdomain formation in a different cellular compartment or if it only functions inside of peroxisomes we attached it to the C-terminus of a mitochondrial reporter protein containing an N-terminal mitochondrial targeting signal. This chimeric fusion protein was detected inside of mitochondria in distinct foci suggesting that the TIIV peptide triggers subdomain formation independent of the targeted organelle (Fig. 3G).

### Identification of additional proteins enriched in the core structure

Intrigued by the identification of a short peptide motif that triggers concentration of GFP inside of intraperoxisomal foci we asked whether this could help uncovering additional protein residents of the peroxisomal core structure.

We mined the predicted *U. maydis* proteome for candidates, which contain a TIIV motif or a similar sequence in combination with a PTS1 (Tab. S1). We tested seven full-length candidate proteins fused to GFP. Localization of six candidates resembled the so far identified proteins Mac1, Mac3 and Uox1. The GFP signal accumulated within peroxisomal foci and in organelle subpopulations (Fig. 4A and 4C; Fig. S4A). This strengthens our notion that the TIIV-like motifs are involved in concentrating proteins into the core structure. Proteins from a control set of randomly chosen PTS1-containing proteins that lack a TIIV-like motif on average showed a higher co-localization with mCherry-SKL (Fig. 4B and 4C, Fig. S4B). The variability of correlation coefficients, however, suggests that other features in the proteins contribute focal enrichment in the core structure and formation of subpopulations as well.

**Figure 4.**
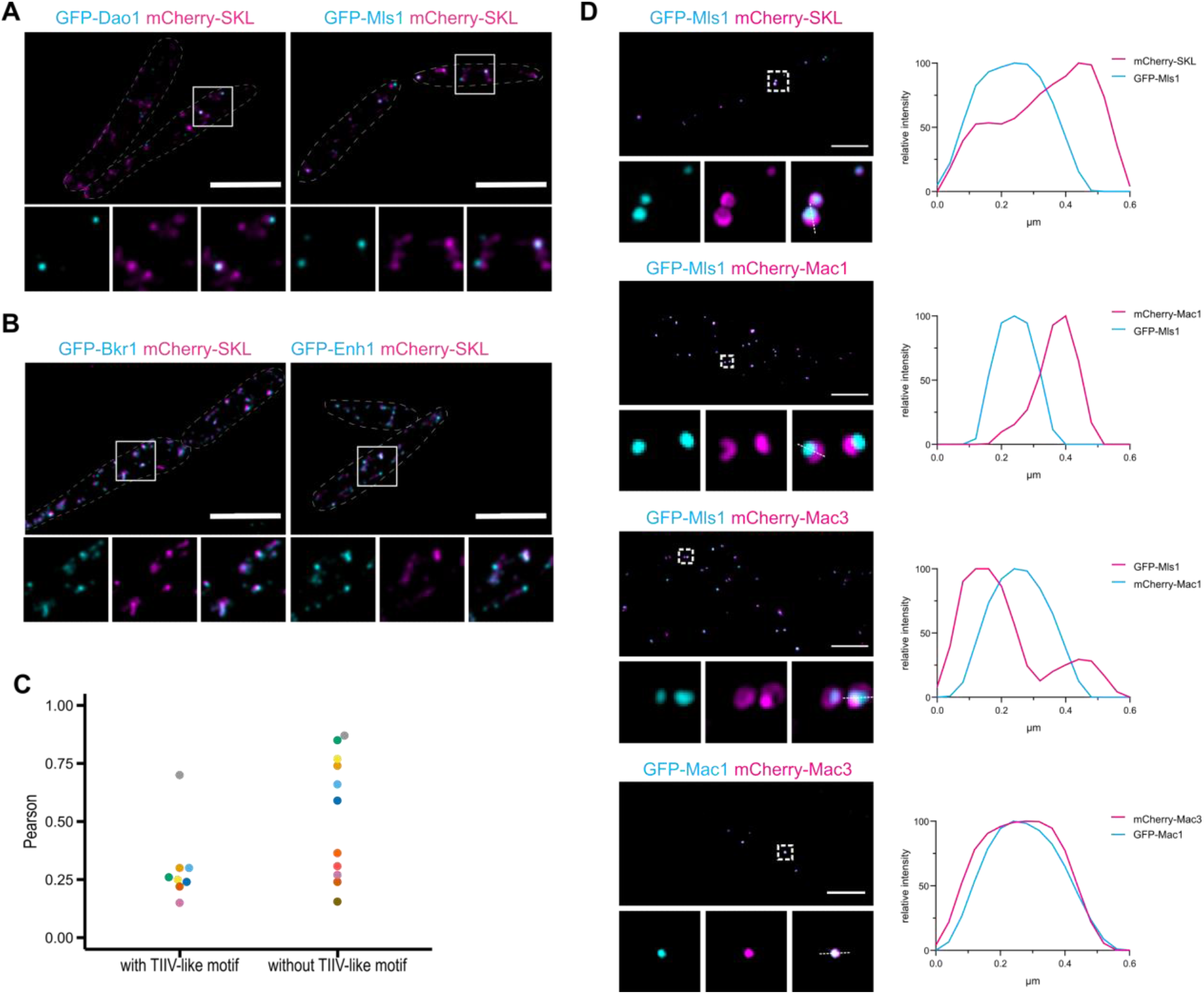
Identification of additional PTS1 proteins enriched in the peroxisomal core structure. (A) Two candidate proteins containing TIIV-like motifs – D-amino oxidase 1 (Dao1) and malate synthase 1 (Mls1) – were co-expressed with mCherry-SKL. Representative epifluorescence images are shown. Full images are provided as overlays of the green and red channel. For insets single channels and merged channels are depicted. (B) Two candidate proteins without TIIV-like motif – a putative NADPH dependent beta-ketoacyl reductase (Bkr1) and a putative enoyl-CoA hydratase (Enh1) – were co-expressed with mCherry-SKL. Representative epifluorescence images are shown. (C) Quantification shows the mean value of Pearson’s correlation coefficients of GFP signals (candidate proteins) and mCherry-Signal signals (mCherry-SKL). Quantifications for three replicates for each of the analyzed candidate proteins are shown in Fig. S4. (D) Co-localization of indicated fusion protein was analyzed by SIM. Organization of pictures as above. Associated graphs show the normalized fluorescence intensity of GFP and mCherry along the indicated lines. (A, B and D) Signals of GFP-tagged proteins are shown in cyan and of mCherry-tagged proteins in magenta. Scale bars: 5 μm.

Together, these data demonstrate that a significant number of peroxisomal matrix proteins accumulates in peroxisomal foci. Other proteins have no or minor affinity to these structures or are unable to form them (Fig. 4). The variable ability of proteins to form or accumulate in subdomains likely represents a basic regulatory principle, which determines the luminal distribution of peroxisomal matrix proteins. Among the enriched proteins were enzymes involved in different metabolic pathways. These include the predicted D-amino oxidase Dao1 (UMAG_05703) and the malate synthase Mls1 (UMAG_15004) involved in glyoxylate metabolism (Fig. 4A). Both proteins are also typical residents of peroxisomes in other organisms (Kunze and Hartig, 2013; Pollegioni et al., 2007), but are not necessarily connected in a metabolic network. Accordingly, enrichment of a particular protein does not coincide with a specific peroxisomal pathway, but rather seems to be a feature shared by peroxisomal proteins with very different biochemical functions. Interestingly, although Mac1, Mac3 and Mls1 are detected in overlapping peroxisomal subdomains and enrich in similar subpopulations they appear to be concentrated inside of different regions of the structures and may even exclude each other to some extent (Fig. 4D), suggesting internal heterogeneity and a complex intrinsic organization of the core structures.

### Peroxisome subpopulations form over time

Enrichment of proteins in peroxisomal subdomains could result in their asymmetric distribution when peroxisomes divide. This may support the formation of peroxisome subpopulations. To follow GFP-Mac3 localization over time, GFP-Mac3 was expressed under control of an arabinose inducible and glucose repressible promoter (Brachmann et al., 2001). As a control, GFP fused to acyl-CoA oxidase 1 (Aox1; UMAG_02208) was used. GFP-Aox1 was not enriched inside of the core, but uniformly distributed among peroxisomes upon constitutive expression (Fig. S5A). Induction of both GFP fusion proteins (GFP-Mac3 or GFP-Aox1) in strains that also express mCherry-SKL, was accomplished through incubation in arabinose containing medium (Fig. 5A).

**Figure 5.**
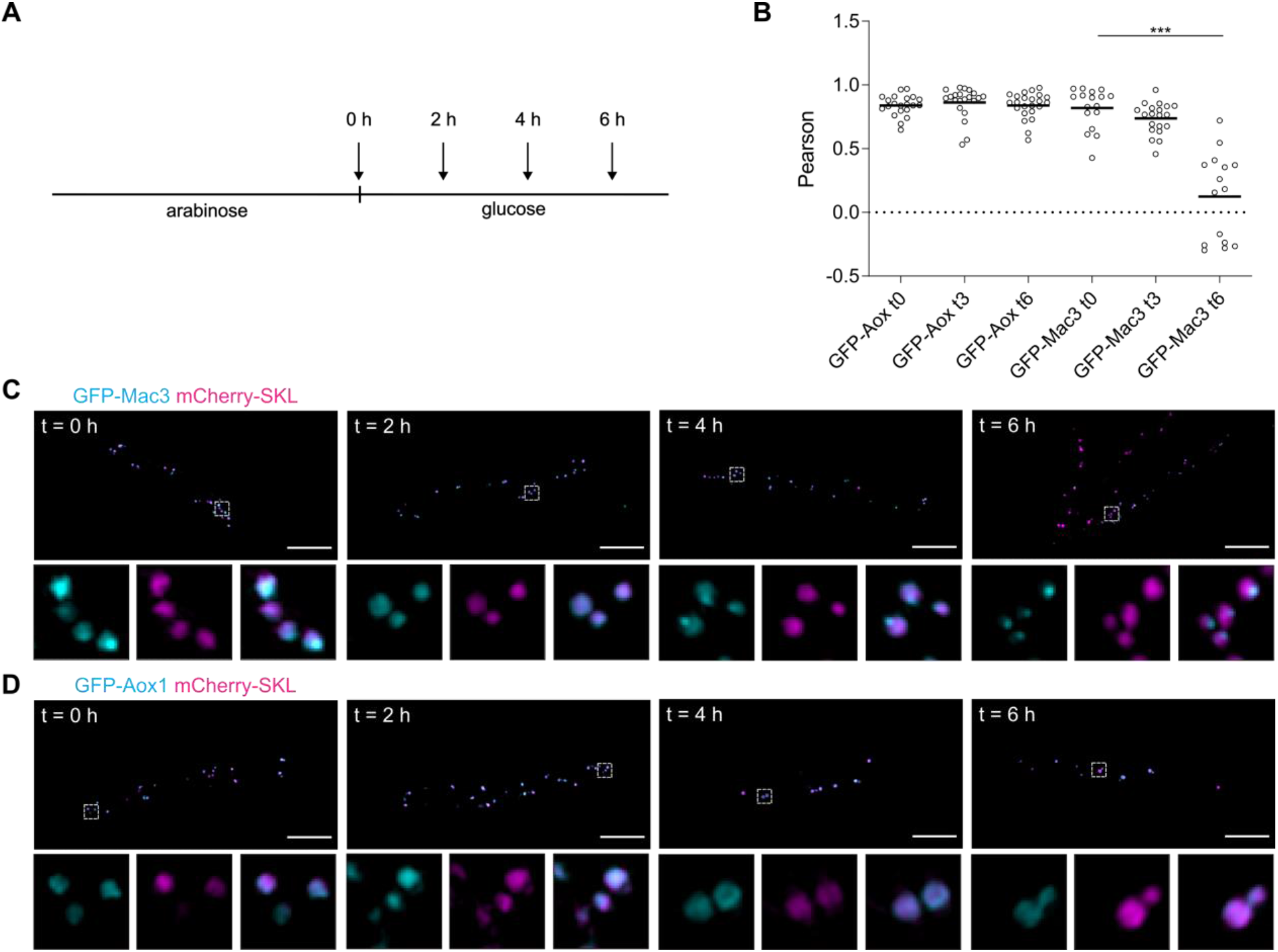
Accumulation in core structures and formation of subpopulations over time. (A) Scheme of the transcriptional pulse-chase experiment to assess the formation of subdomains and subpopulations over time. Cells were incubated in arabinose medium for 2 hours to induce expression of GFP-tagged proteins. Expression was stopped via change to glucose containing medium. (B) Quantification of colocalization data derived from epifluorescence microscopy depicted in Fig. S5. Each circle represents one cell. (C and D). Cells were analyzed at indicated time points by super-resolution microscopy using SIM. GFP-Mac3 (C; cyan) was compared to GFP-Aox1 (D; cyan) – a protein not accumulating in the core structure upon constitutive expression (Fig. S5) – in strains containing mCherry-SKL (magenta). Full images are shown as overlays of the green and red channel. For insets single channels and merged channels are depicted.

After repression of gene expression by shifting cells to glucose medium we followed the localization of the fluorescent proteins at different time points by SIM and epifluorescence microscopy (Fig. 5A to 5D). Quantifications of epifluorescence images revealed that GFP-Mac3 containing subpopulations form over time (Fig. 5B and S5B). Both GFP-Mac3 and GFP-Aox1 were first uniformly distributed among peroxisomes (Fig. 5C and 5D). At later time points, GFP-Mac3, but not GFP-Aox1, accumulated in distinct foci within peroxisomes demonstrating formation of subdomains (Fig. 5C and 5D).

### Specialization of peroxisome populations

We reasoned that the propensity of matrix proteins to self-assemble may control the protein content of specialized peroxisomes. This is reminiscent to the biogenesis of peroxisome derived Woronin bodies in filamentous ascomycetes (Jedd and Chua, 2000).

These specialized peroxisome derived compartments are enriched in a protein dubbed Hex1 in *N. crassa*, which forms crystalline structures resistant to detergent (Jedd and Chua, 2000; Liu et al., 2011, 2008; Yuan et al., 2003). We hypothesized that subdomain formation in *U. maydis* and Woronin body formation represent special adaptations of a more generic mechanism that regulates formation of distinct peroxisome populations. We, therefore, fused GFP to the *Aspergillus nidulans* Woronin body protein HexA (Momany et al., 2002; Zekert et al., 2010) and co-expressed GFP-HexA in *U. maydis* together with either mCherry-Mac3 or mCherry-SKL. mCherry-Mac3 and GFP-HexA were detected in highly overlapping foci whereas mCherry-SKL and GFP-HexA showed a lower degree of co-localization (Figs. 6A and 6B). mCherry-Mac3 and GFP-HexA also appeared in stable and overlapping structures following treatment of crude organelle extracts with detergent (Fig. 6C). This data supports the concept that formation of luminal peroxisomal subdomains in *U. maydis* and formation of Woronin bodies in ascomycetes follows similar mechanistic principles.

**Figure 6.**
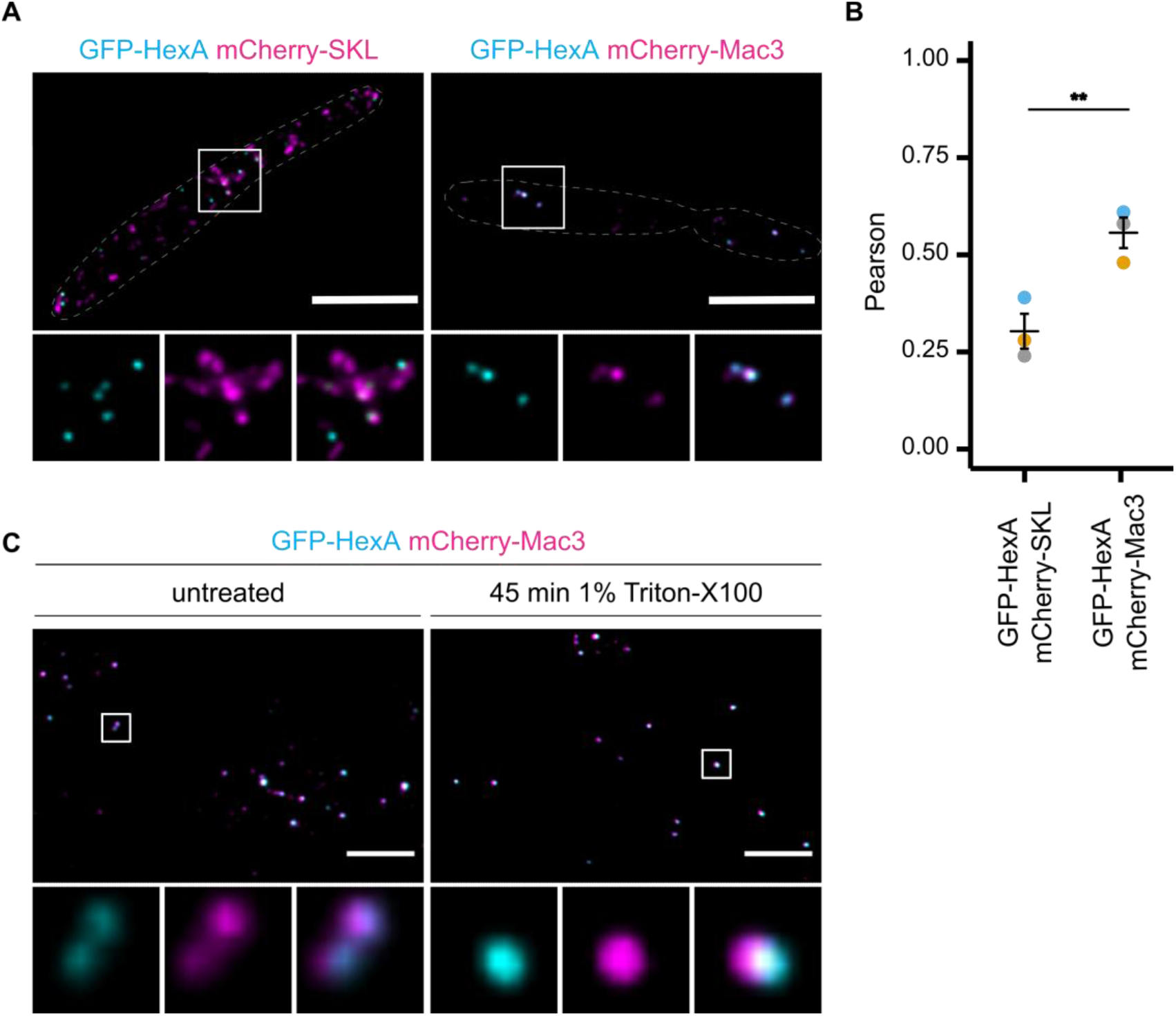
The Woronin body protein HexA colocalizes with Mac3. (A) *U. maydis* strains expressing an N-terminally GFP-tagged version of *A. nidulans* HexA (cyan) and mCherry-SKL (magenta) or mCherry-Mac3 (magenta) were inspected by epifluorescence microscopy. Full images are shown as overlays of the green and red channel. For insets single channels and merged channels are depicted. (B) Quantification shows Pearson’s correlation coefficients of GFP and mCherry signals for indicated strains. (C) Crude organelle preparations of indicated strains were imaged by epifluorescence microscopy after incubation in lysis buffer supplemented with Triton X-100 (right). Preparations incubated in lysis buffer without Triton X-100 served as control (left). Organization of pictures as described above.

### TIIV dependent accumulation in peroxisomal subdomains of mammalian cells

Finally, we addressed transferability of our finding to a different biological system and expressed GFP combined with a minimal PTS1 ending on SKL and containing the TIIV motif in human retinal pigment epithelial 1 (RPE1) cells.

mCherry-SKL was again used as marker for peroxisomes and subcellular localization of fluorescent proteins was analyzed by SIM. We obtained similar data as in *U. maydis* and observed accumulation of GFP-TIIV-SKL in defined subdomains of peroxisomes in RPE1 cells (Fig. 7A). To test if these structures are also enriched for a protein found in the native paracrystalline core of mammalian peroxisomes we co-expressed GFP-TIIV-SKL together with mCherry fused to murine urate oxidase 1 (*Mm*UOX1). GFP-TIIV-PTS1 showed a profound co-localization with mCherry-*Mm*UOX1 revealing that both proteins are located in vicinity (Fig. 7B). Thus, the rules underlying protein concentration inside of the paracrystalline core are likely similar in fungi and in humans. To further confirm this theory we performed complementary experiments in *U. maydis* cells and expressed GFP fused to murine *Mm*UOX1 together with either mCherry-SKL or mCherry-Mac3. Again, we detected a higher degree of overlap for GFP-UOX1 and mCherry-Mac3 than for GFP-*Mm*UOX1 and mCherry-SKL (Fig. 7C). In addition, we could observe stability of GFP-*Mm*UOX1 containing structures upon detergent treatment of crude organelle extracts (Fig. 7D). Both results are consistent with our model, that in diverse eukaryotic species formation of the core structure of peroxisomes follows analogous principles and is a major driving force to spatially organize the peroxisomal proteome.

**Figure 7.**
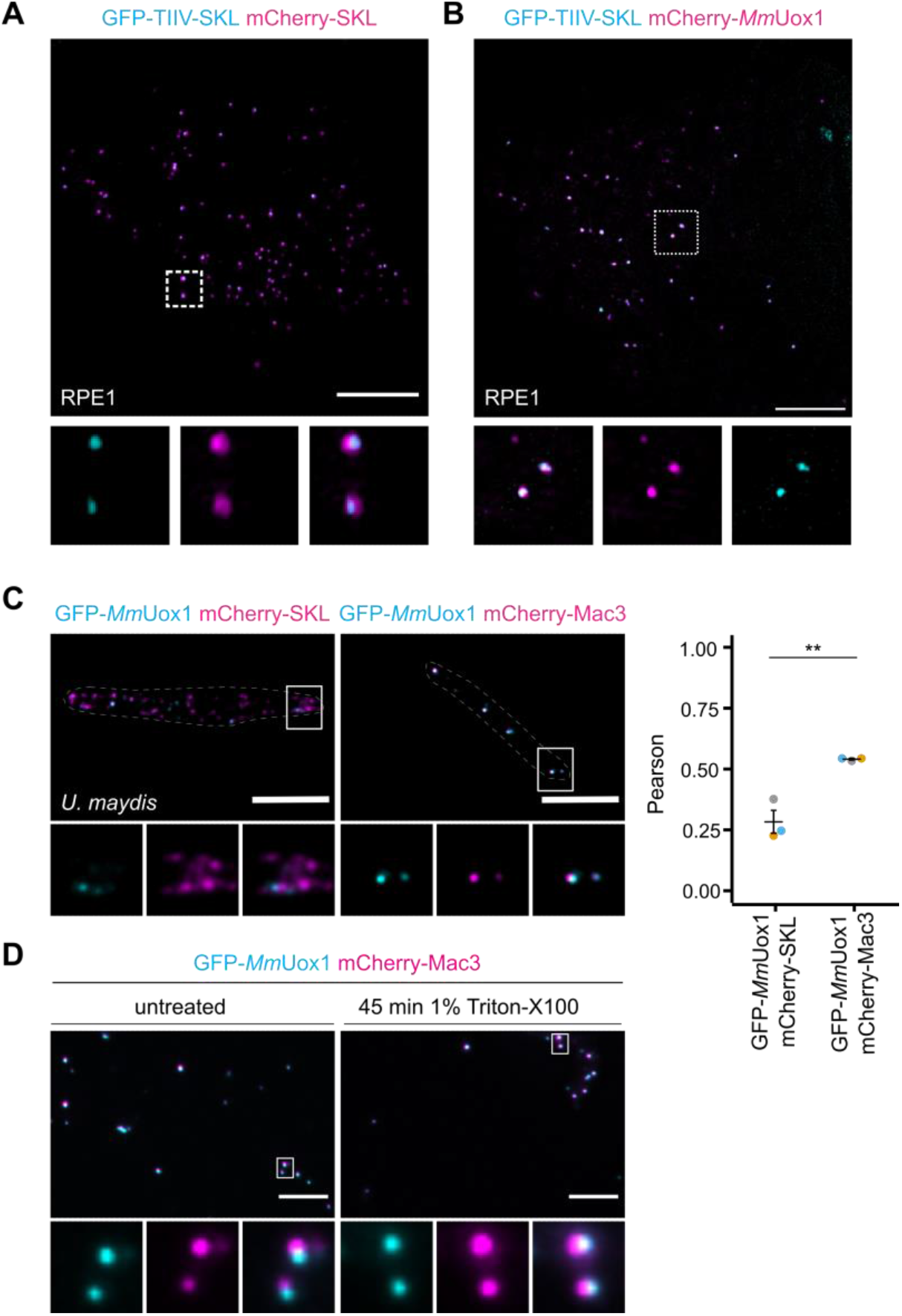
Similar mechanistic principles for enrichment in the core of *U. maydis* and mammalian peroxisomes. (A) Constructs expressing GFP-TIIV-SKL and mCherry-SKL were transfected into RPE1 cells. Cells were analyzed by SIM. The representative picture shows an overlay of the GFP-signal (cyan) and the mCherry-signal (magenta) in the overview. The signal of single channels as well as the overlay are depicted for magnified insets. (B) *Mus musculus* urate oxidase (*Mm*UOX1) was tagged with mCherry (magenta) and co-expressed with GFP-TIIV-SKL (cyan) in RPE1 cells. Cells were analyzed by SIM. Representative pictures are organized as above. (C) *M. musculus* urate oxidase (*Mm*UOX1) was tagged with GFP (cyan) and expressed in *U. maydis* cells either co-expressing mCherry-SKL (magenta) or mCherry-Mac3 (magenta). Representative pictures are organized as above. (D) Quantifications show Pearson’s correlation coefficients of GFP and mCherry signals for indicated strains. (E) Crude organelle preparations of indicated strains were imaged by epifluorescence microscopy after incubation in lysis buffer supplemented with Triton X-100 (right). Preparations incubated in lysis buffer without Triton X-100 served as control (left). Organization of pictures as described above. Scale bars: 5 μm.

## Discussion

Here, we have shown selective accumulation of various proteins in subdomains of peroxisomes likely identical to the paracrystalline core structure. This process turned out as a driving force for the generation of organelle subpopulations with different protein content. Remarkably, independent of the original organism or biochemical function all so far tested proteins enriched in these structures show a high degree of co-localization in *U. maydis* and in human cells (Figs. 1 – 4, Fig. 7). Even the very specialized protein HexA from *A. nidulans* exhibited a very similar pattern of localization suggesting that selective enrichment of this peculiar peroxisomal protein required for Woronin body biogenesis (Jedd & Chua, 2000) follows parallel mechanistic principles (Fig. 6).

The variable ability to self-assemble inside of the lumen of the peroxisome may be a fundamental principle to determine the proteome of every single peroxisome. Peroxisomes proliferate by fission, constantly divide but in contrast to other organelles may not regularly fuse at least in their mature form (Islinger et al., 2012). Thus, the asymmetric segregation of the core will lead to peroxisome subpopulations with different protein content after any fission event. Combined with altered protein expression levels this will almost certainly derive many unique peroxisome variants within a single cell. Furthermore, peroxisomes can accumulate huge amounts of cargo protein (DeLoache et al., 2016; Grewal et al., 2021) and the ability of proteins to accumulate in the core structure may be one prerequisite or even a catalyzer. Peroxisomes are attractive organelles for implantation of synthetic pathways (DeLoache & Dueber, 2013; Simon et al., 2020; Stehlik et al., 2014). Forced concentration of enzymes by addition of TIIV-like motifs might be another way to optimize or to modulate these pathways. Protein crystallization for structural determination purposes (Schönherr et al., 2018) might benefit from our finding as well.

Our data provide first insights into a novel code, which will evolve our understanding of the spatial organization of the peroxisomal proteome. The identification of sequence motifs sufficient for focal accumulation is a starting point to systematically deduce structural differences between proteins that tend to exhibit a homogenous luminal peroxisomal distribution or reside in peroxisomal subdomains. The core structure is, most probably, metabolically active as it is a major site of a number of abundant enzymes including catalase, urate oxidase or alcohol oxidase of methylotrophic yeasts (Hayashi et al., 1976; Veenhuis et al., 1981; Völkl et al., 1988; Heinze et al., 2000; Schönherr et al., 2018). Given the surprising variability of proteins with the ability to enrich in this structure (Fig. 3), its internal organization is of considerable interest. Can proteins go in and out regularly? Are their different layers of enrichment for particular proteins as indicated by our data for acyltransferases and malate synthase 1 from *U. maydis* (Fig. 4)? Is there biochemical activity everywhere in the core? The process of formation may even exhibit functional parallels with phase separation of proteins (Boeynaems et al., 2018; Hyman et al., 2014).

In summary, our work contributes to a more detailed characterization of a conserved peroxisomal structure, which is known since the discovery of the peroxisome (De Duve & Baudhuin, 1966; Rouiller & Bernhard, 1956) but whose formation and composition has not been addressed in greater detail until now.

## Methods

### Microorganism, growth conditions and transformation

*Escherichia coli* strain TOP10 (Invitrogen) was used for all cloning purposes and amplification of plasmid DNA. *U. maydis* strains generated and used in this study are derivatives of Bub8, MB215 and Bub8 mCherry-SKL (Freitag et al., 2012; Hewald et al., 2005; Weinzierl et al., 2002) and listed in Supplementary Table 2 (Tab. S2). All strains are available from the authors upon reasonable request. *U. maydis* cells were grown at 28°C in YEPSlight medium (Tsukuda et al., 1988) or yeast nitrogen base medium (YNB; Difco) supplemented either with 2% glucose and 0.2% ammonium sulfate or 0.2% oleic acid (Roth), 0.1% Tween-40 and 0.2% ammonium sulfate at pH 5.6. For epifluorescence imaging cells were grown in oleic acid media if not specified differently. SIM imaging was performed on glucose grown cells to circumvent artifact of oleic acid addition. For chase experiments (Fig. 5) *U. maydis* cells were grown overnight in YEPSlight medium. Subsequently, cells were washed and diluted in YNB medium containing 2% arabinose and 0.2% ammonium sulfate and incubated for 2 h. Expression of fusion proteins was stopped by shifting the cells to YNB medium containing 2% glucose and 0.2 % ammonium sulfate cells and the localization of the fluorescent proteins was microscopically monitored at indicated time points. Cell transformation was performed as previously described (Hanahan et al., 1991; Schulz et al., 1990). DNA was integrated into the *ip*-locus of *U. maydis* cells (Broomfield & Hargreaves, 1992) or randomly integrated into the genome. Hygromycin (200 μg/ml), carboxin (2 μg/ml) and G418 (400 μg/ml) were used for selection. Genomic DNA was extracted as previously described (Hoffman and Winston, 1987).

### Human cell line and transient transfection of cells

hTert-RPE1 cells (ATCC CRL-4000) were cultured in DMEM-F12 medium (Life Technologies) supplemented with 10% FCS and Penicillin/Streptomycin (100 U/mL penicillin, 100 μg/mL streptomycin) at 37°C, 5% CO2. Cells were transiently transfected with the help of Fugene 6 (Promega) using a ratio of 3:1 of reagent to DNA. Cells were transfected in MatTek 3.5 cm glass-bottom dishes. 12 h after transfection, the medium was replaced to remove transfection reagent, and the cells were imaged between 16 h and 20 h post transfection.

### Molecular cloning and nucleic acid procedures

Standard procedures were followed for plasmid generation (Sambrook et al., 1989). Plasmids used for protein expression in *U. maydis* are derivatives of the plasmids pOTEF-GFP-Ala6-MMXN and pCRG-GFP-Ala6-MXN (Böhmer et al., 2008). Plasmids used for expression of fluorescent proteins in hTert-RPE1 cells are derivatives of the plasmid pEGFP-C1 and pmCherry-C1 (Clontech). Plasmids were analyzed by sequencing. All plasmids and oligonucleotides used in this study are listed in Supplementary table S2 (Tab. S2). Plasmids are available from the authors upon reasonable request.

### Epifluorescence microscopy

Logarithmically grown *U. maydis* cells were treated with 100 μM Carbonyl cyanide m-chlorophenyl hydrazine (CCCP) resolved in dimethyl sulfoxide to block peroxisomal movement. For the majority of experiments using epifluorescence microscopy cells were incubated in oleic acid-containing media for four hours prior to imaging experiments as formation of subpopulations was easier to follow under these conditions (exceptions: imaging experiments underlying Figs. 1A and S5). After 5-10 minutes incubation at room temperature, the cells were either placed on 1% agarose cushions containing 100 μM Carbonyl cyanide m-chlorophenyl hydrazine CCCP on a microscope glass slide and covered with a coverslip or placed on MatTek 3.5 cm glas-bottom dishes and covered with an 1% agarose cushion containing 100 μM CCCP as well. For time-lapse imaging no CCCP was added. Epifluorescence microscopy was performed on an Axiovert 200 M inverse microscope (Zeiss) equipped with a 1394 ORCA ERA CCD camera (Hamamatsu Photonics), filter sets for eGFP and rhodamine, and a Zeiss 63x Plan Apochromat oil lens (NA 1.4). The software Volocity 5.3 (Perkin-Elmer) was used for image acquisition. The ImageJ plugin DeconvolutionLab with 25 iterations of the Richardson-Lucy algorithm was used for image deconvolution (Vonesch and Unser, 2008), point-spread functions for each channel were computed by the ImageJ plugin PSF Generator using the Richards & Wolf model (Kirshner et al., 2013). Alternatively, epifluorescence microscopy as well as time-lapse experiments (500 ms time-lapse for 60 s) were performed on a Deltavision OMX V4 microscope (GE Healthcare) equipped with three water-cooled PCO edges CMOS cameras, a solid-state light source and a laser-based autofocus. The software softWoRx (Applied Precision, GE Healthcare) was used for image deconvolution. Images were processed in ImageJ/Fiji (Schneider et al., 2012).

### SIM

Three-dimensional structured illumination microscopy (SIM) was performed on a Deltavision OMX V4 microscope with and Olympus x60 NA 1.42 objective and three PCO.edge sCMOS cameras and 488 nm and 568 nm laser lines. Cells were illuminated with a grid pattern. 15 raw images (3 orientations, 5 phases) were taken for each image plane. SoftWoRx software (Applied Precision, GE Healthcare) was used for image reconstruction, alignment and projection. Images were processed in ImageJ/Fiji (Schneider et al., 2012). For 3D visualization of peroxisomal subdomains, surfaces were generated using Imaris.

### Transmission electron microscopy and Immunogold labelling

For visualization of *U. maydis* cells expressing GFP-Mac3 using transmission electron microscopy (TEM) 50 ml logarithmically growing cells were harvested and frozen under high-pressure (Wohlwend HPF Compact 02). After subsequent freeze substitution (using acetone, containing 0.25% osmium tetroxide, 0.2% uranyl acetate, 0.05% ruthenium red, and 5% water) (Leica AFS2), cells were embedded in Epon812 substitute resin (Fluka). Embedded cells were sectioned to 50 nm thin sections (Leica EM UC7 RT), which were used for immunolabeling with α-GFP (Rockland; dilution 1:500). As a secondary antibody rabbit-anti-goat antibodies coupled to ultra-small gold particles were used (Aurion, dilution 1:100). Subsequently a silver enhancement procedure was performed and sections were post-stained with 2% uranyl acetate and 0.5% lead citrate. Analysis of the samples was conducted with a JEOL JEM2100 TEM equipped with a fast-scan 2k CCD TVIPS (Gauting, Germany) F214 camera. Images were processed with ImageJ (Schneider et al., 2012).

### Preparation of crude organelles

Preparation of post-nuclear supernatants was adapted from previous protocols (Cramer et al., 2015; Kremp et al., 2020; Stehlik et al., 2020). Briefly, 200 ml of logarithmically growing cells were harvested for 5 min at 1600 x g at 23°C. Cells were washed two times with deionized water and two times with sorbitol buffer (1.2 M sorbitol; 20 mM KH_2_PO_4_ adjusted to pH 7.4 with K_2_HPO_4_). Formation of spheroplasts was achieved through incubation in sorbitol buffer containing novozyme (4μg/ml) for 30 – 45 min. Degradation of the cell wall was followed microscopically. Afterwards, spheroplasts were kept on ice and gently washed two times with sorbitol buffer – centrifugation for 10 min at 2300 rpm at 4°C. Spheroplasts were gently washed with lysis buffer (5 mM MES, 0.5 mM EDTA, 1 mM KCl, 0.6 M Sorbitol, 1 mM 4-aminobenzamidine dihydrochloride, 1 μg/ml aprotinin, 1 μg/ml leupeptin, 1 mM phenylmethylsulfonyl fluoride, 10 μg/ml N-tosyl-L-phenylalanine chloromethyl ketone, and 1 μg/ml pepstatin) – centrifugation for 10 min at 2300 rpm at 4°C, resuspended in 15 ml lysis buffer and frozen at −80°C overnight. Homogenization was achieved through 2 × 10 strokes with a Potter-Elvehjem homogenizer interrupted by chilling the samples on ice for 2 min. Nuclei and cell debris were removed by two subsequent centrifugations at 1600 × g for 10 min. Subsequently, the PNS was diluted to an OD600 of 1, aliquoted and frozen at −80°C.

### Detergent treatment

PNS fractions were thawed on ice, centrifuged at 10k*g for 10 min at 4°C and concentrated 10x in lysis buffer. Samples were split into halves and aliquots were incubated in lysis buffer containing 1% Triton-X-100 or in lysis buffer for 45 min. Preparations were immediately analyzed by epifluorescence microscopy immobilized on 1% agarose cushions.

### Quantification of peroxisomal subpopulations and statistical analysis

Microscopic data was collected from at least three independent *U. maydis* cultures (biological replicates) for each experiment. At least five images per culture with at least ten cells were quantified. Quantification of co-localization was performed as follows. Detection and measurement of individual peroxisomes was performed automatically via an ImageJ macro (Supplementary File 1). Briefly, maximum projections of deconvolved RFP and GFP z-stacks were created, transformed into 16-bit images and corrected for channel-misalignment by the “StackReg 2.0.0” plugin using “Rigid Body” transformation (Thevenaz et al., 1998; https://sites.imagej.net/BIG-EPFL/plugins/) to later serve as the reference images for detection of peroxisomes; same operations were performed on average projections of the respective non-deconvolved z-stacks for later fluorescence-intensity measurements. To increase sensitivity and account for peroxisomes showing only fluorescence in one of the two channels, the resulting images were merged using the Image Calculator function with the maximum option (detection based on only RFP or GFP references was still included in the macro but performed less reliable than using the merge of both references). For detection of peroxisomes, the merged average projections were used to create a mask of the cells. This mask was used to determine the background noise outside of the cells in the reference merge. The resulting maximum noise intensity was used for the Find Maxima function of ImageJ; individual peroxisomes were detected in the reference merge and marked as point selections. These selections were restored in the background-corrected average projections of the non-deconvolved stacks of both channels. Subsequently, circular ROIs of a 3-pixel radius (area of 32 pixels) were created around the point selections and signal intensities were measured for both channels (as Raw Integrated Density per ROI) and stored side-by-side in the results file for the whole dataset. A commented version of the macro is attached to the manuscript. For evaluation of the resulting data, the statistics software R was used (R Core Team, 2015): measured intensities were normalized to one and a linear regression analysis was performed with the RFP intensities as independent and GFP intensities as dependent variable, in addition a correlation test was performed (Pearson’s). Superplots (Lord et al., 2020) and statistical tests were computed using RStudio 1.2.1335 with R 3.6.0 or GraphPadPrism. Plots are structured as follows: center line, mean; error bars, standard error of the mean; circles, mean of experiments. P-values were calculated using an unpaired, two-sided Student’s t-test using Graphpad Prism. For data depicted in Fig. 1 and Fig. S3C, which contain multiple comparisons a 1-Way anova combined with a Tukeys post-test was performed to assess significance of the differences. * refers to a p-value lower than or equal to 0.05; ** refer to a p-value lower than or equal to 0.01 and ***lower than or equal to 0.001. ****lower than 0.0001. The graph shown in Fig. 5 was generated using Graphpad Prism. Each dot represents one cell.

### Bioinformatics and accession numbers

*U. maydis* putative peroxisomal PTS1 protein sequences were retrieved from the *Ustilago maydis* genome database (MUMDB) or broad institute and manipulated with notepad ++ using regular expressions (regex). PTS1 motifs were bookmarked with the regular expression ‘([SA][RKHQNS][LI]|[SA][RK][MFV]|[PCVGE][RK][LI])\*$‘. Protein sequences containing a TIIV-like motif were bookmarked with the regular expression ‘(T [I, L, V] [I, L, V] [I, L, V] [I, L, V] [I, L, V])\*$’. Peroxisomal candidate proteins with TIIV-like motif and without motif were randomly chosen and the localization of the peroxisomal proteins to subdomains was analyzed by fluorescence microscopy. All genes can now be accessed via the National Center of Biotechnology Information (NCBI) via the UMAG number.

## Data availability

Any supporting original data for this study can be requested from the corresponding authors.

## Acknowledgments

We thank Marisa Piscator and Ulrikke Dahl Brinch for excellent technical assistance. We are grateful to Reinhard Fischer and Valentin Wernet for plasmids. JA was supported by a fellowship from Marburg Research Academy. TS acknowledges funding from SYNMIKRO. KS was supported by a Career grant from the South-Eastern Norway Regional Health Authority (2020038) and a Research Grant from the Research Council of Norway (315103). JF acknowledges funding from the DFG (grant ID FR-3586/2-1).

## Author contributions

Conceptualization: JA, DM, KS and JF; Data acquisition: JA, NB, DM, TH, BS, KS and JF. Preparation of figures: JA, NB, DM and KS; Generation of code, bioinformatic and statistical analysis: JA, NB, TS, CR, KS; Supervision: CT, GB, MB, KS, JF; Writing – original draft: JF; Writing – revision and editing: All authors. Analysis of data: All authors.

## Supplementary figures

**Figure S1.**
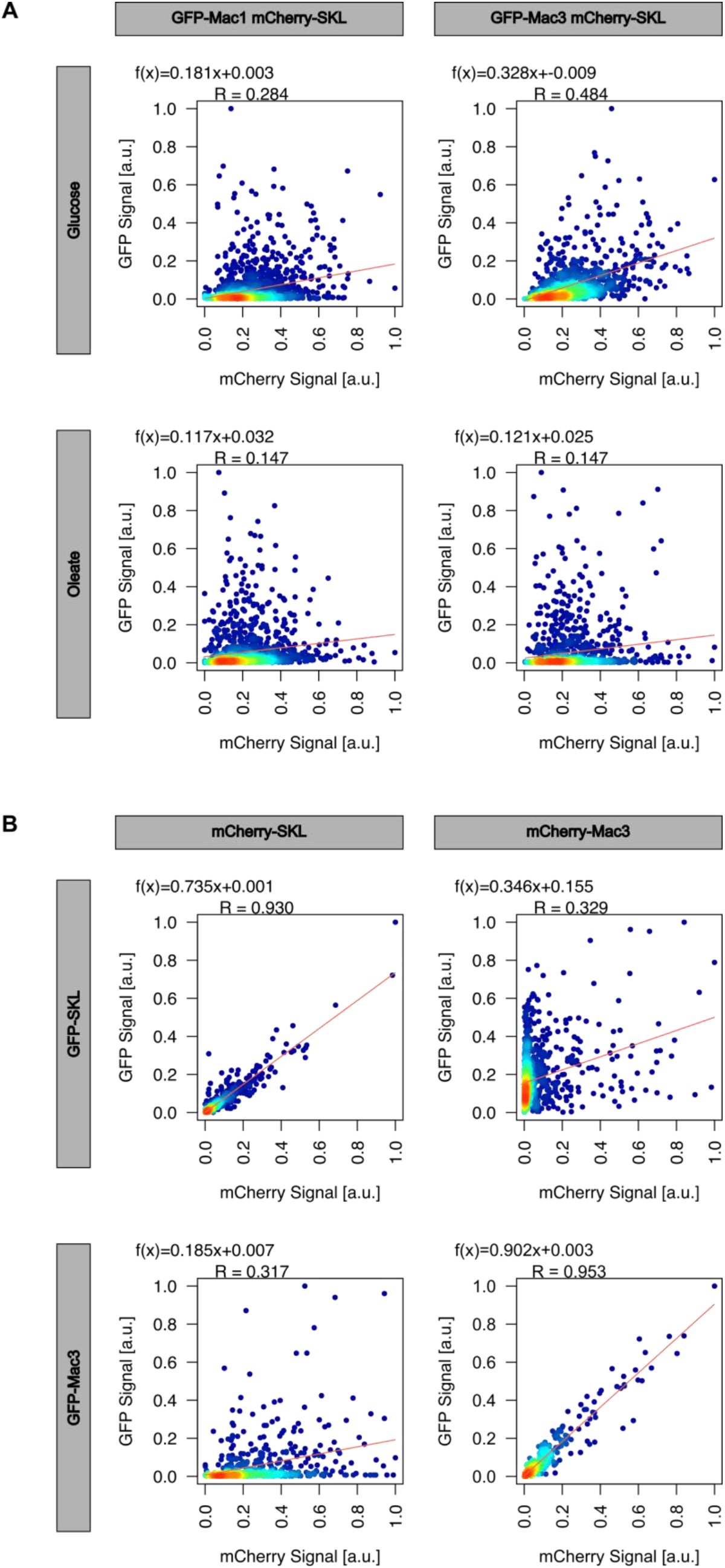
Examples for the quantification of co-localization. Representative plots showing the fluorescence signals of all analyzed peroxisomes in a representative replicate. (A) related to Fig. 1C (B) related to Fig. 1F.

**Figure S2.**
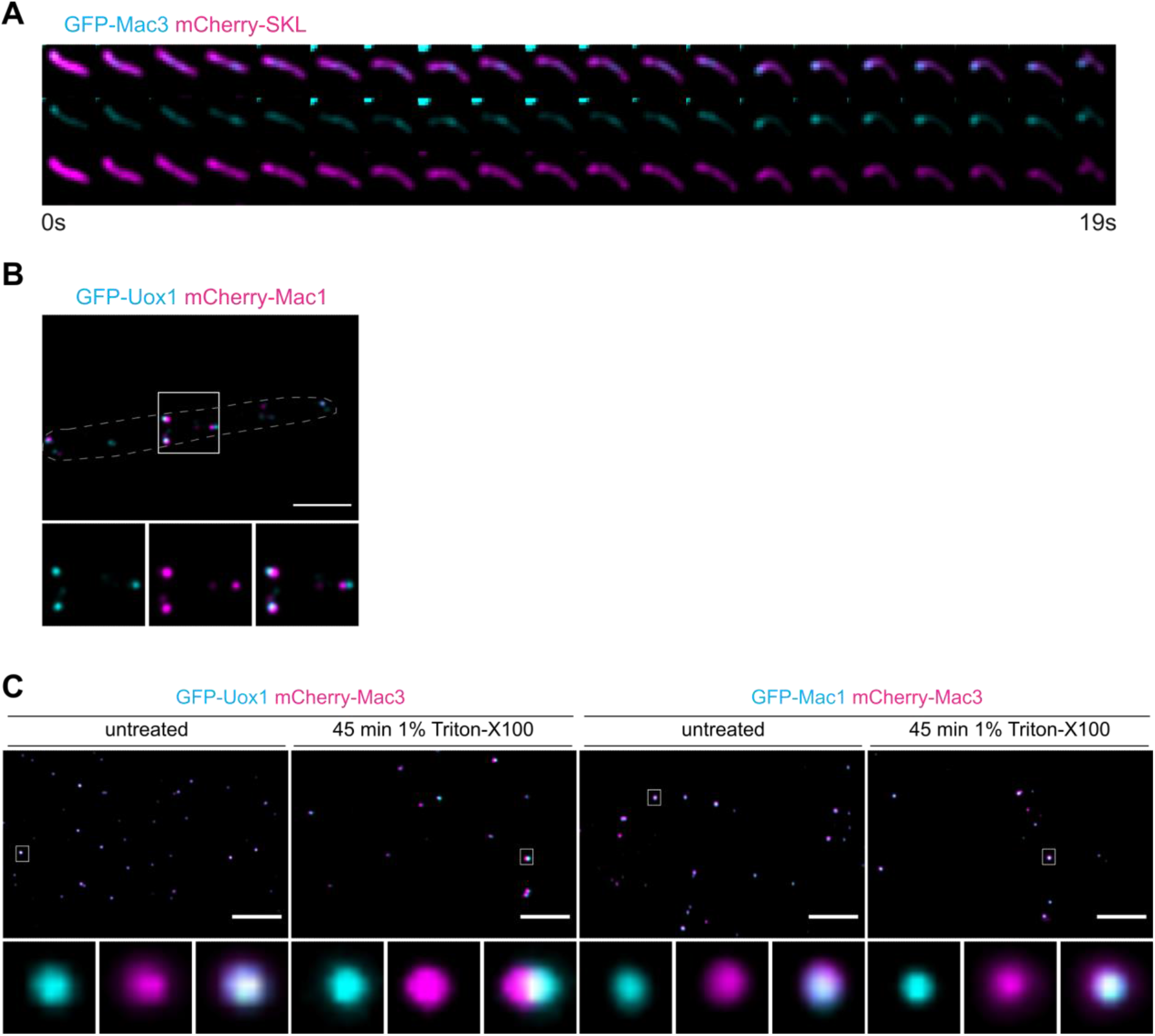
Mac proteins localize in detergent resistant peroxisomal subdomains also enriched for urate oxidase 1. (A) Time lapse imaging related to Movie 1. Epifluorescence imaging of GFP-Mac3 (cyan) and mCherry-SKL (magenta). One frame per second was recorded. Shown are overlays of the mCherry and GFP signal. (B) A representative image of cells expressing GFP-Uox1 and mCherry-Mac1. Full images are shown as overlays of the green and red channel. For insets single channels and merged channels are depicted. Scale bar: 5 μm. (C) Crude organelle preparations of indicated strains were imaged by epifluorescence microscopy after incubation in lysis buffer supplemented with Triton X-100 (right). Preparations incubated in lysis buffer without Triton X-100 served as control (left). Organization of pictures as described above. Scale bars: 5 μm.

**Figure S3.**
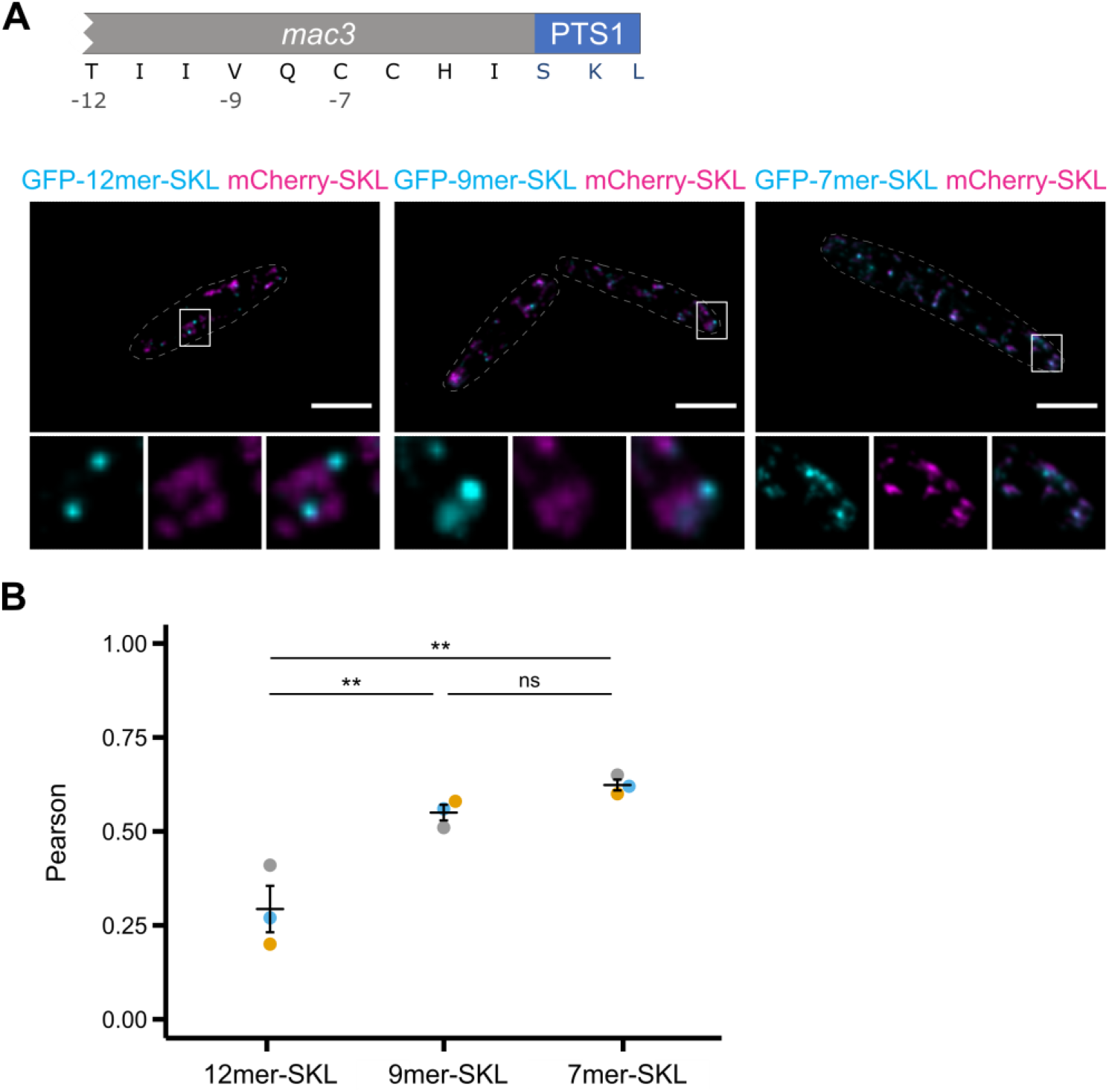
Truncation analysis of the C-terminal dodecamer of Mac3. (A) Depiction of the 3’end of the *mac3* gene encoding the peroxisomal targeting signal PTS1. Please note that the unusual ASL motif had to be substituted by SKL to perform the subsequent truncation analysis as truncated fragment of the dodecamer did not allow full peroxisomal import, which perturbs co-localization analysis (B) Indicated fragments of the C-terminal dodecamer were fused to GFP (cyan) and analyzed in strains also expressing mCherry-SKL (magenta). Full representative images are shown as overlays of the green and red channel. For insets single channels and merged channels are depicted. Scale bars: 5 μm. (C) Quantification of co-localization of experiments shown in (B).

**Figure S4.**
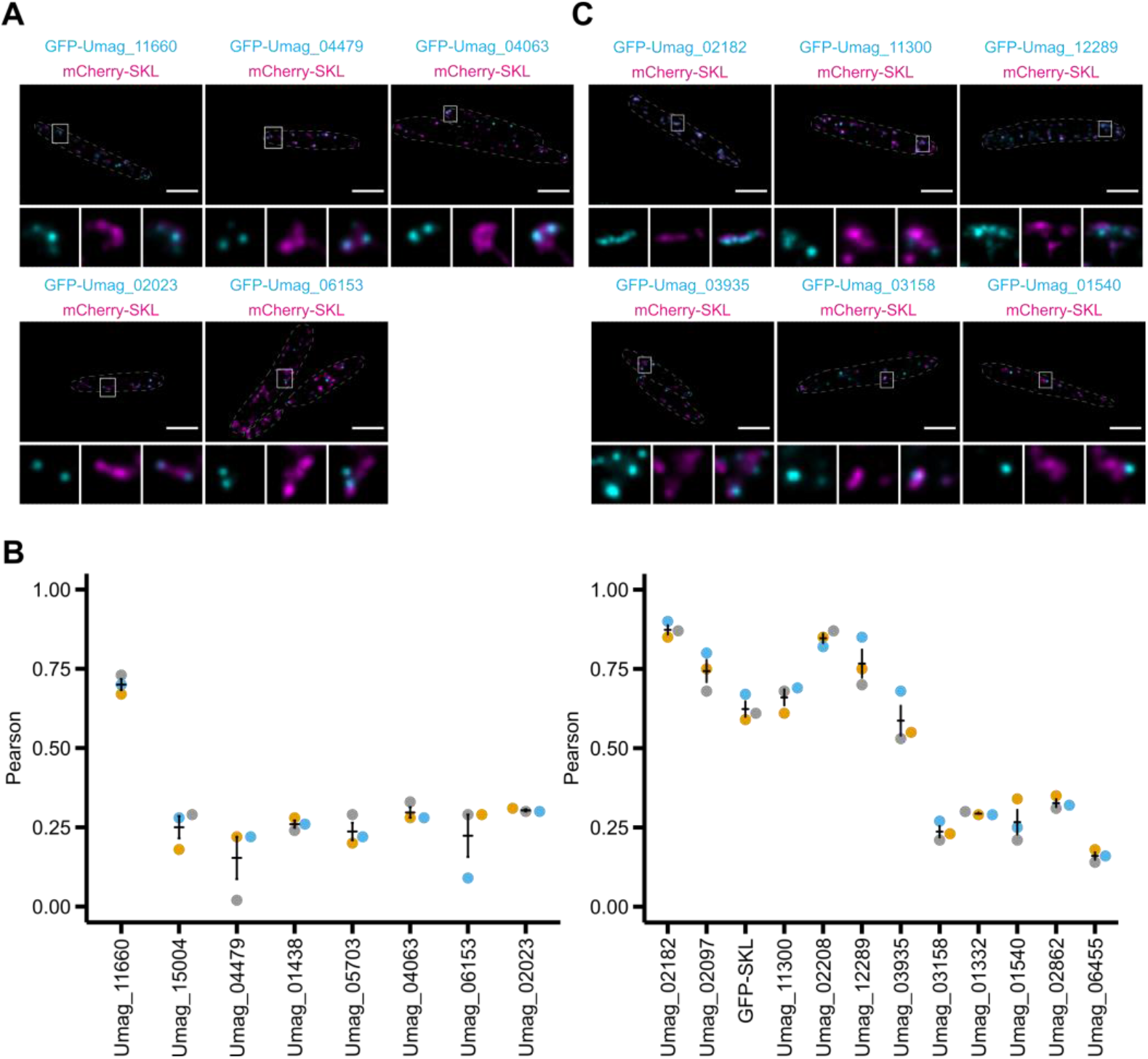
Analysis of candidate proteins with or without TIIV-like motif. *U. maydis* strains expressing N-terminally GFP-tagged versions of candidate proteins (cyan) with TIIV like motifs (A) or without TIIV-like motifs (C) and the peroxisomal marker protein mCherry-SKL (magenta) were inspected by epifluorescence microscopy. Full representative images are shown as overlays of the green and red channel. For insets single channels and merged channels are depicted. Scale bars: 5 μm. Umag numbers are gene identifiers accessible via NCBI. (B) Quantifications show Pearson’s correlation coefficients of GFP and mCherry signals for indicated strains.

**Figure S5.**
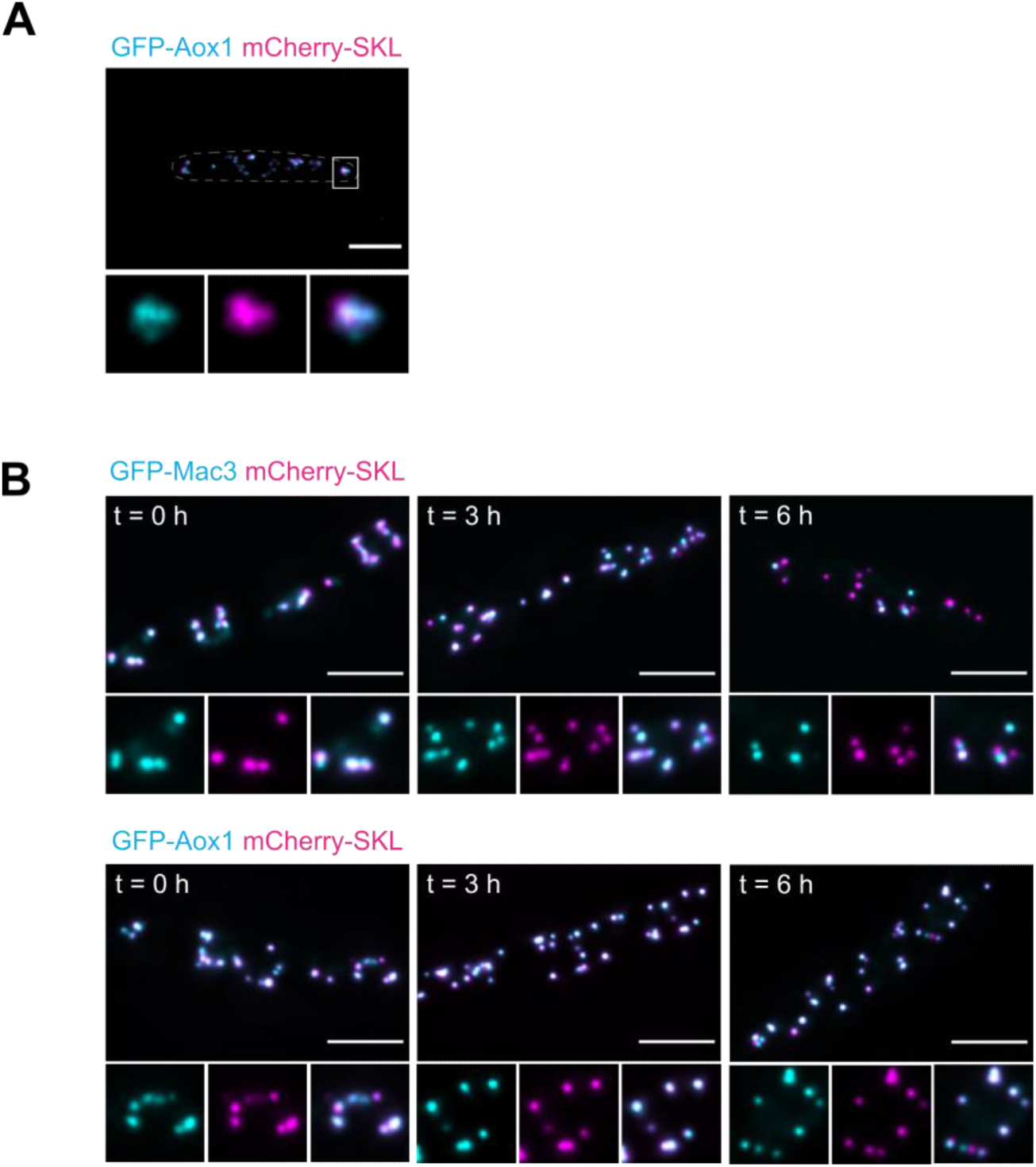
Formation of peroxisome subpopulations over time. (A) *U. maydis* strains expressing an N-terminally GFP-tagged version of Aox1 (cyan) and the peroxisomal marker protein mCherry-SKL (magenta) were inspected by epifluorescence microscopy. Full images are shown as overlays of the green and red channel. For insets single channels and merged channels are depicted. Scale bar: 5 μm. **(B)** Cells were analyzed at indicated time points after addition of glucose by epifluorescence microscopy. GFP-Mac3 (cyan) was compared to GFP-Aox1 (cyan) in strains containing mCherry-SKL (magenta). Full images are shown as overlays of the green and red channel. For insets single channels and merged channels are depicted. Quantifications of (B) are shown in Fig. 5.

## Notes

### Competing Interest Statement

The authors have declared no competing interest.

### Summary of Updates

This manuscript has been revised to correct a mistake unfortunately introduced in the amino acid code.

